# Neutrophil Extracellular Traps Mediate *In Vitro* and *In Vivo* Degradation of α-Synuclein Amyloid Fibrils

**DOI:** 10.1101/2025.09.26.678871

**Authors:** Thyago R. Cardim-Pires, Ariele da Silva Martins, Fernanda Verdini Guimarães, Thayana Roberta Ferreira de Mattos, Elvira Saraiva, Patrícia Machado Rodrigues Silva, Debora Foguel

## Abstract

Neutrophil extracellular traps (NETs) are chromatin-derived structures decorated with neutrophil enzymes such as elastase and myeloperoxidase. Our group has previously demonstrated that amyloid fibrils (AFs), regardless of their protein composition, induce NET release *in vitro* in human neutrophils through a process dependent on reactive oxygen species (ROS) generation by NADPH oxidase 2 (NOX-2). Moreover, the proteases embedded in NETs were shown to degrade AFs into smaller, potentially toxic species. The present study aimed to determine whether amyloid fibrils composed of α-synuclein (αSF) can induce NET formation *in vivo* and to investigate the role of NETs in modulating amyloid-associated pathology. To this end, we employed gp91phox knockout (KO) mice, which lack NOX-2 activity and are therefore unable to release NETs. αSF was instilled into the lungs of both WT and KO mice (males and females), leveraging the lung’s robust immune cell recruitment - particularly of neutrophils-as a model system. Eight hours after αSF instillation, both WT and KO animals exhibited marked neutrophil infiltration in the lungs causing inflammation. However, NET formation-evidenced by the presence of citrullinated histones and myeloperoxidase - was detected only in WT mice. Interestingly, while Congo red-positive amyloid-like structures persisted in the lungs of KO mice, they were absent in the lungs of WT animals, suggesting that NET-associated proteases facilitate the clearance of AFs from lung tissue. Lung function was assessed by measuring elastance and resistance. Our data showed that, while AFs were still present in the lungs of both WT and KO mice, elastance was impaired. As AFs were cleared from the lungs of WT mice, lung function recovered. In contrast, KO animals, in which AFs persisted, continued to exhibit compromised elastance. Together, our findings demonstrate that AFs impair lung function, and that NETs, induced in response to these fibrils, promote their degradation and thereby protect lung tissue from further damage.

## Introduction

Neurodegenerative diseases such as Parkinson’s disease (PD) and Alzheimer’s disease (AD) can be classified as forms of amyloidosis, as they share a common hallmark: the accumulation of protein aggregate species, including amyloid fibrils (AFs) and oligomers. Each condition is distinguished by the specific protein from which the aggregates originate and by their anatomical distribution across the brain and peripheral tissues^1,2^. A wide range of peptides and proteins can adopt this fibrillar conformation, including the Aβ peptide and α-synuclein (αSyn), which are primarily associated with AD and PD, respectively^3–5^. Regardless of the protein of origin, aggregation follows a conserved pathway in which proteins undergo conformational changes that first give rise to small toxic oligomers during the nucleation phase, which then progress into protofibrils and ultimately mature AFs through exponential growth^1,6,7^. Importantly, oligomers are not merely transient intermediates but are actively involved in a variety of toxic cellular processes, including the formation of membrane pores that promote cytotoxicity, synaptic dysfunction, endoplasmic reticulum stress, and impaired autophagy, among others^7,8^.

The presence of protein aggregates triggers inflammatory responses and promotes the release of cytokines, chemokines, and other immune mediators that recruit and activate effector cells^9–12^. Neutrophils, the most abundant leukocytes in the blood, are among the first responders to tissue injury and play a crucial role in host defense^13^, a role sustained by their highly diverse antimicrobial arsenal stored within distinct granule subsets^14,15^. Key defense mechanisms include degranulation, phagocytosis, and the formation of neutrophil extracellular traps (NETs), a specialized response involving the release of DNA-histone complexes together with granular proteins into the extracellular space^14,16^.

NETs are chromatin-decondensed structures released by neutrophils and are decorated with a wide array of different proteins, including elastase, myeloperoxidase, as well as antimicrobial peptides, other bioactive molecules^17,18^. This highly conserved immune response has been observed across multiple species^19^ and was initially described as a mechanism for trapping and killing invading microorganisms^16^. Nowadays, the involvement of NETs has been recognized far beyond their antimicrobial function. Excessive or dysregulated NET formation, as well as impaired clearance, has been linked to tissue damage and to the pathogenesis of a wide range of inflammatory and autoimmune diseases^20–22^.

Depending on the stimulus, NET formation can proceed through either ROS-dependent or ROS-independent mechanisms^20^. In the ROS-dependent pathway, NET release is primarily driven by superoxide (O_2_^−^) production catalyzed by the NADPH oxidase complex (NOX-2). NOX-2 is a transmembrane enzyme complex whose catalytic subunit, gp91phox, reduces molecular oxygen to superoxide by oxidizing NADPH^23,24^. A broad range of stimuli can trigger NET release by elevating ROS levels, including pathogens of different origins (viruses, bacteria, fungi), activated platelets, pro-inflammatory cytokines, and even AFs^17,24–28^.

Our group was the first to demonstrate that human neutrophils stimulated with AFs of three distinct proteins - αSyn, transthyretin (TTR), and Sup35 - release extracellular traps *in vitro* in a NOX-2 dependent manner^27,28^. Additionally, we found that elastase, a major component of NETs, is capable of degrading TTR-AFs into toxic oligomers. We also observed that NET markers, including elastase and extracellular DNA, colocalized with amyloid deposits in lungs and skin biopsies from patients with systemic amyloidosis, supporting the relevance of this interaction.

Building on previous evidence, our study investigated whether NETs induced by AFs composed of αSyn could contribute to amyloid pathology *in vivo* by instilling AF in the lungs of wild-type (WT) and gp91phox-knockout (KO) mice, which lack the catalytic subunit of NOX-2. These animals are a well-established model of chronic granulomatous disease and are widely employed to study NET deficiency in settings where the stimulus is ROS-dependent^29,30^.

Our findings revealed that although both WT and KO mice showed neutrophilic infiltration and inflammation following AF instillation, only WT animals were able to clear αSyn fibrils from the lungs. In addition, fibril clearance in WT animals was accompanied by the restoration of pulmonary function, whereas lung dysfunction persisted in KO mice where fibrils remained. These findings suggest that NETs contribute not only to the inflammatory response but also to the clearance of amyloid material. To our knowledge, this is the first report demonstrating both the deleterious impact of amyloid accumulation in the lungs and the critical role of neutrophil extracellular traps in mediating amyloid degradation (amyloidolysis) and promoting recovery of organ function.

## Materials and Methods

### Ethic Committees’ Approvals

All animal experiments in this study were approved by the Ethics Committee on the Use of Animals (CEUA) of the Health Sciences Center at the Federal University of Rio de Janeiro. Procedures were conducted under process number 01200.001568/2013-8, protocol nº 004/23, in accordance with established ethical guidelines for animal research. All in vitro experiments involving human neutrophils were approved by the Research Ethics Committee (CEP) of the Clementino Fraga Filho University Hospital, Federal University of Rio de Janeiro (HUCFF-UFRJ), under protocol number 4261.015400/0052-57.

### Animal Model

In this study, we used male and female C57BL/6 mice (2–4 months old), including both wild-type and gp91phox knockout animals. gp91phox encodes the catalytic subunit of the NOX-2 complex, and its deletion abolishes the generation of ROS via NOX-2. Because the gp91phox gene is located on the X chromosome, male knockout mice are hemizygous (gp91phox^y/–^), whereas females are homozygous (gp91phox^−/−^). For consistency, all gp91phox-deficient animals are referred to as gp91phox KO mice throughout the manuscript, irrespective of sex. The sex of the animals used in each experiment is indicated in the figure legends and methods sections.

### *In vitro* experiments

#### Purification of Mouse Bone Marrow Neutrophils

Murine neutrophils were isolated from the bone marrow of euthanized mice following exposure to 5% isoflurane. Femurs and tibias were carefully removed and placed in 1× Hank’s balanced salt solution (HBSS). Bone marrow cells were flushed from the cavities using a 1 mm needle with 3 mL of 1× HBSS, dissociated, and centrifuged at 400 × g for 5 min. Erythrocytes were lysed by incubation in 10 mL of 0.2% NaCl for 40 s, after which osmolarity was restored by adding 10 mL of 1.6% NaCl. Cells were centrifuged and resuspended in 5 mL of 1× HBSS before being layered over a two-step Percoll gradient (65% and 72%). Gradients were centrifuged at 1000 × g for 30 min with the brake off. Neutrophils were collected from the 65– 72% interface, washed twice with 1× HBSS to remove residual Percoll, and finally resuspended in RPMI medium. Neutrophil purity was assessed by cytocentrifugation followed by panoptic staining.

#### Assessment of NET Production via Extracellular DNA Detection

To evaluate NET production, purified neutrophils (5 × 10^5^ cells/well) from WT and gp91phox KO animals were seeded in a 96-well plate (Costar®), and stimulated with 10 μM αSyn fibril (αSF), 100 nM PMA (Sigma-Aldrich - P8139) or 1μg/mL of LPS O111:B4 (Sigma-Aldrich, #L3023) for 3 h. NETs were measured using Sytox Green (Invitrogen, Cat. No. S7020) according to the manufacturer’s instructions and imaged in a EVOS M5000 Imaging System (Thermo Fisher Scientific). For experiments with hydrogen peroxide (H_2_O_2_), it was pre-diluted at 1 μM in RPMI medium and added to the cells accordingly.

#### Determination of Reactive Oxygen Species (ROS) Production

To assess ROS production, 5 × 10^5^ neutrophils from WT or gp91phox KO mice were incubated with 100 nM phorbol myristate acetate (PMA, positive control), 10 μM αSF, and 1.2 μM dihydrorhodamine 123 (DHR, Sigma-Aldrich). Fluorescence intensity was measured using a SpectraMax microplate reader (Molecular Devices), with excitation at 500 nm and emission at 540 nm.

#### Production of Supernatant of human PMA-activated Neutrophil (SPAN)

Purified neutrophils from humans (8 × 10^6^ cells/well) were cultured in 24-well plates (Corning) and incubated with 100 nM PMA at 37 °C in 5% CO_2_ for 4 h. The supernatants were then collected and centrifuged at 400 × g for 10 min. The pellet was discarded, and the supernatant was stored at -80 °C until further use. NET quantification was performed using the PicoGreen dsDNA kit (Invitrogen Life Technologies) according to the manufacturer’s instructions.

#### In Vitro Digestion of Amyloid Fibrils by Neutrophil Elastase and SPAN

To evaluate the digestion of αSF by neutrophil elastase and SPAN, 5 μM of αSF were incubated with 10 μg/mL NE (Sigma-Aldrich, #324681) and 1 μg/mL SPAN at 37°C for 4 h. Aliquots were withdrawn at 0, 15, 30, 60, 90, 120, 180, and 240 min, and αSF degradation was monitored by 15% SDS-PAGE followed by Western Blotting using a PVDF membrane previously activated with methanol. The membrane was blocked with Intercept (PBS) Blocking Buffer (LI-COR^®^) for 1 h, incubated with anti-human α-Syn antibody (1:5,000; Thermo Fisher Scientific) overnight, and with the corresponding secondary antibody (IRDye^®^ 800CW, 1:5,000, LI-COR^®^) for 2 h. Visualization was performed on the LI-COR^®^ Odyssey Scanner 9120, and quantification was done on ImageJ software (National Institutes of Health). Fibril digestion was further evaluated by Transmission Electron Microscopy (TEM) using negative contrast with 2% uranyl and by Thioflavin T binding using the ChronosDFD ISS spectrofluorimeter (ISS Inc., Champaign, IL, USA) with excitation at 450 nm and emission from 470 nm to 600 nm.

#### Expression and Purification of Human Wild-Type α-Synuclein

Purification of α-Synuclein was performed as described by ^31^. To do this, human wildtype αSyn was expressed in *Escherichia coli* BL21 (DE3) pLysS cells transformed via heat shock with the corresponding plasmid. Protein expression was induced with 0.5 mM IPTG (isopropyl β-Dthiogalactopyranoside, Sigma-Aldrich) and cultures incubated for 4 h under the same conditions. Cells were harvested (6000 rpm, 15 min, 4 °C; Himac CR 21G2, Hitachi), resuspended in 20 mL lysis buffer (20 mM Tris, 5 mM EDTA, pH 8.0) with protease inhibitor cocktail (Thermo Fisher Scientific), and lysed by sonication in ice for 1 h (59 s on/59 s off, 60% amplitude; Vibra Cell™, Sonics). The lysate was centrifuged (16,000 rpm, 20 min, 4 °C), and the pH of the supernatant was lowered to 3.5 with HCl, followed by a second centrifugation and readjustment to pH 7.4 with NaOH. Protein was precipitated with 2 mM ammonium sulfate, and the pellet dialyzed overnight against 20 mM Tris, 1 mM EDTA (pH 8.0), then in Milli-Q water for 4 h before lyophilization. The lyophilized protein was resuspended in PBS (pH 7.4), filtered (0.22 μm), and quantified at 280 nm (NanoDrop® 2000, Thermo Fisher Scientific; ε = 5,900 M^−1^·cm^−1^). Residual lipopolysaccharide was removed using Pierce™ High-Capacity Endotoxin Removal Spin Columns (Thermo Fisher Scientific) according to the manufacturer’s instructions.

#### αSynuclein aggregation and formation of pre-formed fibrils

To induce fibril formation, monomeric αSyn was incubated under constant agitation at 800 rpm for 7 days at 37 °C. The presence of AFs was confirmed by light scattering (excitation and emission at 330 nm), ThT binding assay (10 μM of αSyn; 50 μM of ThT; Excitation at 450 nm and emission collected from 470 to 520 nm; maximum emission at 482 nm) both collected in a Jasco FP8200 spectrofluorometer, Jasco Corp., Tokyo, Japan, and transmission electron microscopy (negative staining, 1% uranyl as contrast agent). The characterization of αSF is shown in **Supplementary Figure 1**.

### *In Vivo* Experiments

#### Instillation of Amyloid Fibrils in the Lung

The animals were anesthetized by aerosolization of isoflurane (4%) and instilled intranasally with 50 μg of α-synuclein fiber diluted in 40 μL of sterile saline (NaCl 0.9%). Control animals received the same volume of saline. After instillation, the animals were subjected to tests to analyze lung function and leukocyte infiltrate.

#### Invasive assessment of respiratory mechanics

Animals were anesthetized via intraperitoneal (i.p.) injection with ketamine (20 mg/kg) and xylazine (10 mg/kg), followed by a single i.p. dose of sodium pentobarbital (60 mg/kg) diluted in distilled water. Tracheostomized animals were mechanically ventilated, and lung function was assessed. The trachea was cannulated and the cannula connected to a pneumotachograph. Airflow and transpulmonary pressure were recorded with a FinePointe R/C Buxco Platform (DSITM, Minneapolis), which calculated resistance (cm H_2_O.s/ml) and dynamic lung compliance (mL/cm H_2_O) in each breath cycle. Elastance was calculated as the inverse of compliance values. Animals were allowed to stabilize for 5 min and increasing concentrations of methacholine (9 - 162 mg/ml) were aerosolized for 5 min each. Baseline pulmonary parameters were assessed with aerosolized PBS.

#### Bronchoalveolar lavage and leukocyte analysis

Leukocytes in the airways were assessed by bronchoalveolar lavage fluid (BALF) collection. Briefly, a tracheal incision was made, and a polyethylene catheter (1.7 mm OD) was inserted. The airways were washed three times with 0.5 mL of phosphate-buffered saline (PBS) containing 10 mM EDTA. BALF was centrifuged (300 × g, 10 min, 4 °C), and cell pellets were resuspended in 250 μL PBS with 10 mM EDTA. For total leukocyte quantification, BALF samples were diluted in Türk solution (2% acetic acid) and cells were counted using a Neubauer chamber under a light microscope (BX40; Olympus, Center Valley, PA). Differential cell counts were performed on cytospin smears stained with May-Grünwald Giemsa and analyzed by light microscopy (BX40; Olympus).

#### Determination of myeloperoxidase (MPO) in the lung

Myeloperoxidase activity was determined using one-third of the tight lung lobe. Cryopreserved lung tissue samples were homogenized (Homogenizer; Omni International, Kennesaw, GA, USA) and centrifuged at 10,000 × g for 10 min. The resulting pellets were resuspended in PBS supplemented with 0.5% HTAB and 5 mM EDTA, followed by centrifugation to isolate the supernatants, which were incubated in a reaction mixture containing PBS-HTAB-EDTA, HBSS, O-dianisidine dihydrochloride (1.25 mg/mL), and 0.4 mM H_2_O_2_. After 15 min at 37 °C with agitation, the reaction was stopped by adding 1% NaN_3_. MPO activity was quantified spectrophotometrically at 460 nm, normalized to total protein content, and expressed as optical density per mg of protein.

#### Histological Analysis

The left lung lobes were removed and fixed in Millonig buffer solution (pH 7.4) with 4% paraformaldehyde, following standard procedures: Dehydration in a graded ethanol series (70% to 100%); 2. Clearing with xylene; 3. Embedding in paraffin; 4. Sectioning using a microtome (5 μm thickness), and 4. Staining with Hematoxylin and Eosin (H&E) for general lung tissue architecture; Picrosirius Red (PS) for collagen deposition, and Congo Red for detection of amyloid fibrils. The slides were coded, and the analyses were carried out blindly. Random airways were photographed using a light microscope Olympus BX40 (Olympus, PA, USA) at x 400 and a digital camera.

#### Assessment of Lung Function Following Amyloid Fibril Instillation

To evaluate lung injury induced by amyloid fibril instillation, animals were pre-anesthetized via intraperitoneal (i.p.) injection with ketamine (20 mg/kg) and xylazine (10 mg/kg), followed by a single i.p. dose of sodium pentobarbital (60 mg/kg) diluted in distilled water. Animals were then positioned on a heated platform within a plethysmograph chamber to maintain normothermia. A tracheostomy was performed, and a cannula was inserted to enable mechanical ventilation, ensuring constant flow and tidal volume parameters. Pulmonary function was assessed using an invasive whole-body barometric plethysmography system (Buxco® System – DSI). Lung elastance (mL/cmH_2_O) and airway resistance (cmH_2_O·s/mL) were measured at baseline and following intratracheal aerosolization of increasing concentrations of methacholine (9–162 mg/mL in PBS). Each dose was delivered over a 5-min period to evaluate airway hyper responsiveness.

#### Immunohistochemistry for NET Detection in Lung Tissue

To assess the presence of NETs in lung tissue, immunohistochemistry was performed following standard deparaffinization and antigen retrieval procedures following protocol described by Ferreira and colleagues (2013) ^32^. Tissue sections were blocked with 5% bovine serum albumin (BSA, Sigma-Aldrich #A7906) and incubated with primary antibodies against myeloperoxidase (1:500; R&D Systems, AF3667) and citrullinated histone H3 (CitH3, 1:250; Abcam AB5103) overnight. Afterward, tissue sections were washed and incubated with fluorophore-conjugated secondary antibodies Alexa Fluor 647 and Alexa Fluor 488 (1:500; Thermo Fisher Scientific A-21447 and A-11008) for 2 h.

## Results

As previously reported, AFs composed of different proteins act as potent ROS-dependent inducers of NETs *in vitro*, using human neutrophils as a model. We also detected the presence of NETs in the lungs and skin of a postmortem patient with primary amyloidosis^27^. Based on these observations, we sought to determine whether NET formation exerts beneficial or detrimental effects in the context of amyloid diseases in an *in vivo* setting.

To address this, we employed a mouse strain deficient in NOX-2-mediated ROS production as an experimental model for NET deficiency. This strain carries a deletion in the gene encoding the catalytic subunit of NOX-2 (gp91phox), which is located on the X chromosome. Consequently, both male (gp91phox^y/-^) and female (gp91phox^-/-^) mice lack functional NOX-2 activity; in the case of females, a double knockout is performed. Both sexes were included in this study, as specified in the experiments.

### AFs induce NET formation in wild-type (WT) murine neutrophils but not in gp91phox knockout (KO) neutrophils *in vitro*

As a first step in our characterization, we sought to determine whether AFs could trigger the release of NETs, a phenomenon previously demonstrated only in human neutrophils. To validate the functional deficiency of NOX-2, we evaluated ROS production in bone marrow neutrophils isolated from gp91phox KO and WT mice following stimulation with PMA, a well-established ROS inducer, or pre-formed α-synuclein fibrils (αSF). As expected, neutrophils from gp91phox KO mice (red symbols) failed to generate ROS in response to both stimuli, whereas WT neutrophils (black symbols) produced significant amounts of ROS, confirming the absence of functional NOX-2 activity in gp91phox KO animals (**Figure 1A** and **inset**).

**Figure 1.**
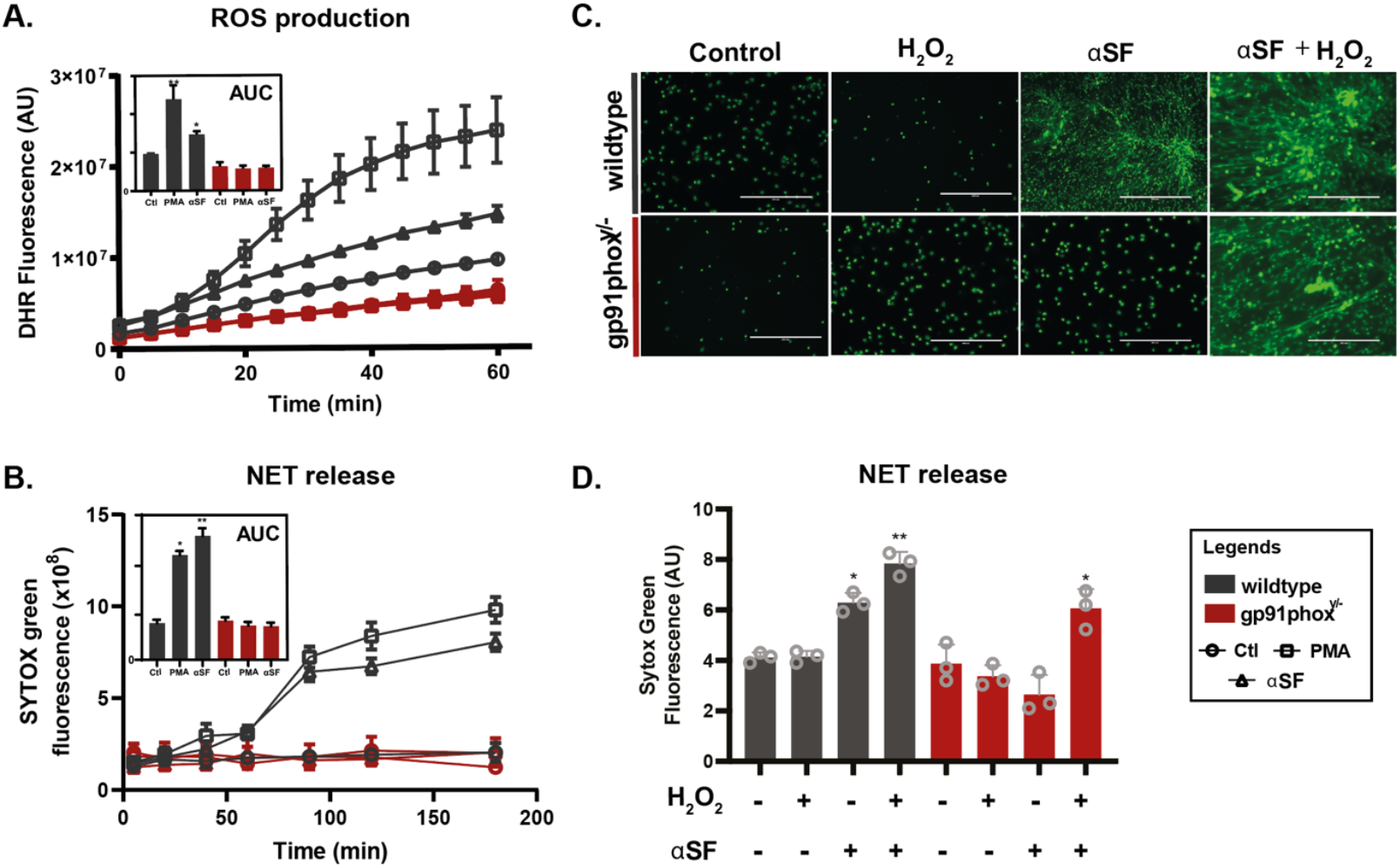
Amyloid fibrils induce neutrophil extracellular trap (NET) formation *in vitro* in a NADPH oxidase-2 (NOX-2) dependent manner. (**A**) ROS production in wild-type (WT) and gp91phox knockout (KO) neutrophils stimulated with PMA or αSF for 1 h. Inset: area under the curve (AUC) analysis. (**B**) Time course of NET release after stimulation with 10 μM αSF or 100 nM of PMA for 3 h. (**C**) Representative images of WT and KO neutrophils stimulated with αSF or αSF + H_2_O_2_ for 2 h. Extracellular DNA was visualized using Sytox Green staining. (**D**) Quantification of NET formation corresponding to the conditions in **panel C**, based on Sytox Green fluorescence Scale bar = 200 μm. Data are expressed as mean ± SEM. *p < 0.05; **p < 0.01 vs. control from 5 independent replicates. Images shown are representative of at least 5 independent replicates. Legends are presented in the right.

Next, we performed a time-course analysis of NET production in WT and gp91phox KO neutrophils stimulated with αSF or PMA. As shown in **Figure 1B**, WT neutrophils (black curves) exhibited increase in Sytox Green fluorescence, an indicative of NET formation, after 3 h of stimulation with αSF or PMA. In contrast, gp91phox KO neutrophils (red symbols) failed to form NETs in response to either stimulus. However, when KO neutrophils were co-stimulated with αSF and a low concentration of H_2_O_2_ (1 μM), they underwent NETosis, as demonstrated by the increase in Sytox Green fluorescence (**Figure 1C** and **D**) and the presence of web-like structures characteristic of NETs visualized by microscopy (**Figure 1C**).

These results demonstrate that murine neutrophils, like human neutrophils, are susceptible to AF-induced NET formation in a NOX-2-dependent manner. gp91phox KO neutrophils fail to generate NETs in response to αSF or PMA unless exogenous H_2_O_2_ is provided, highlighting the critical role of NOX-2-derived ROS in this process.

### αSF triggers the recruitment of immune cells to the lungs independently of NOX-2 activity

Next, to gain better insight into whether AF can trigger the recruitment of immune cells, we instilled αSF into the lungs of both WT and KO mice. The choice of the lung was based not only on its association with systemic amyloidosis, but also on its widespread use in acute inflammation models. As shown in **Figure 2**, both WT and gp91phox KO mice, female (panels **A–F**) and male (panels **G–L**), recruited leukocytes (**A** and **G**), mononuclear cells (**B** and **H**), and neutrophils (**C** and **I**) to their lungs 8 h after αSF instillation, although at different levels. Notably, both sexes of gp91phox KO mice (red bars) displayed a significantly higher number of recruited leukocytes - particularly neutrophils - compared to WT mice (black bars). Interestingly, male mice recruited more neutrophils than female mice, in both WT and gp91phox KO backgrounds. A striking observation was the absence of mononuclear cell recruitment in female WT mice following fibril instillation due to reasons that we do not know at the present.

**Figure 2.**
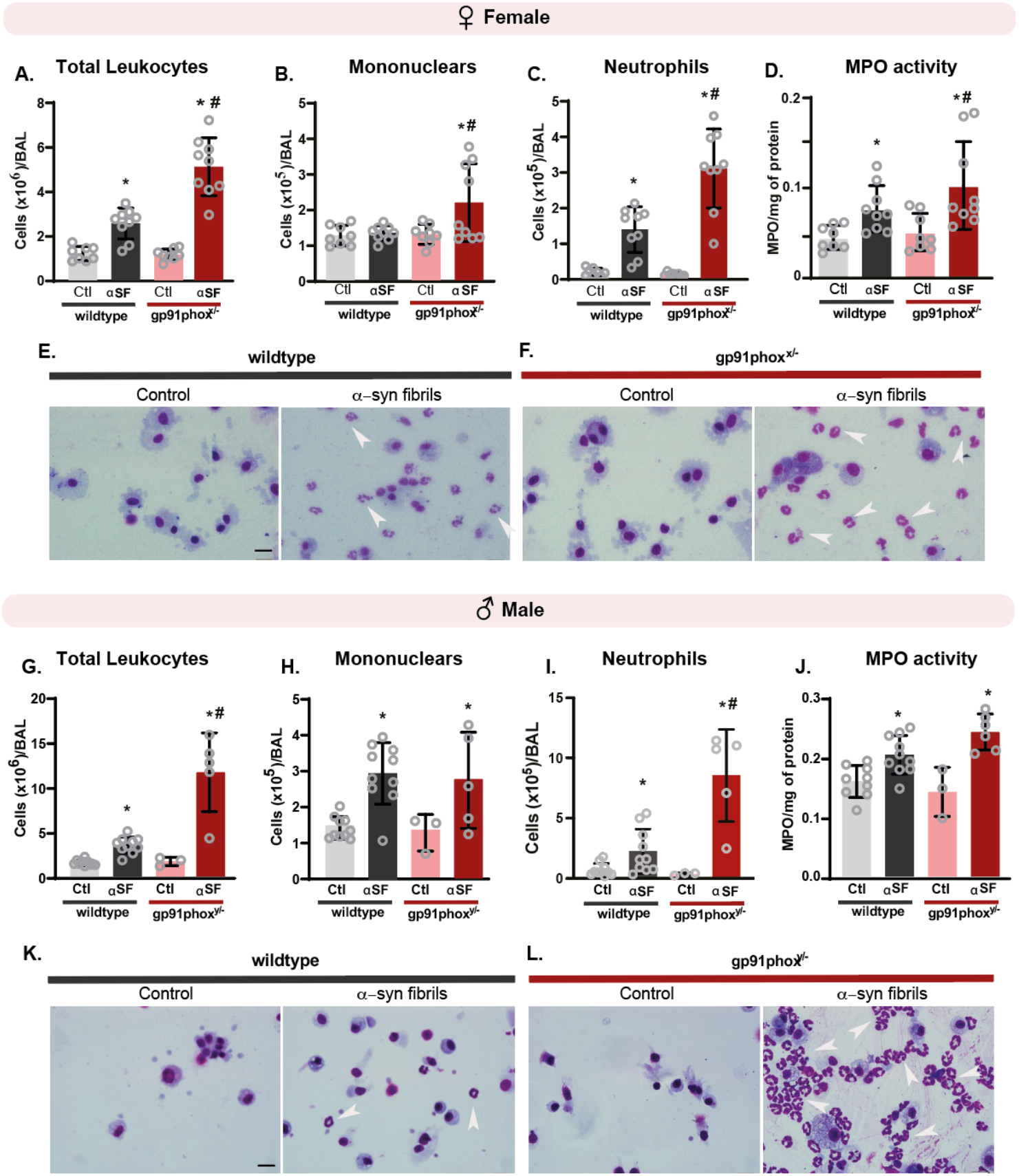
αSyn AFs (αSF) instilled in the lung of WT and gp91phox KO animals recruit leukocytes. 50 μg of αSF were intratracheally instilled in the lungs of the animals and after 8 h cellular recruitment was assessed. (**A** and **G**) Total leucocytes; (**B** and **H**) Mononuclears; (**C** and **I**) Neutrophils and (**D** and **J**) MPO activity were quantified in the BALF. The data are shown separately for females (**A–F**) and males (**G–L**). The lower images show representative BALF images with the recruited immune cells for female (**E** and **F**) and male (**K** and **L**) mice. White arrows point to neutrophils. Scale bar: 100 μm. Results shown as mean ± SEM. * p < 0.05 compared to the control group instilled with saline (Ctl); # p < 0.05 compared to the WT group instilled with αSF. For female mice, sample sizes were n = 8 for controls and n = 9 for αSF-instilled animals. For male mice, wild-type groups included n = 10 for both controls and αSF-instilled, while gp91phox^y/−^ mice included n = 3 for controls and n = 6 for αSF-instilled animals. Scale bar = 100 μm.

Representative images of the bronchoalveolar lavage fluid (BALF) of female and male mice are shown in **panels E** and **F**, and **K** and **L**, respectively. To further support the neutrophil recruitment observed in the BALF, we measured MPO levels in the lungs (**Figure 2D** and **J**). MPO was significantly increased in both gp91phox KO and WT mice (female and male) instilled with AFs compared to saline-treated controls, reinforcing the conclusion that AFs promote neutrophil recruitment.

### Clearance of αSF is impaired in gp91phox KO mice

To further investigate the inflammatory processes induced by αSF instillation in the lungs, we employed a series of histological staining techniques. Hematoxylin and eosin (H&E) staining of lung sections revealed a densely populated pulmonary parenchyma and alveolar infiltration in both WT and gp91phox KO mice (**Figure 3A**). Picrosirius staining further demonstrated collagen deposition following αSF instillation in both genotypes, consistent with an acute inflammatory response. Notably, we observed several “toothpick-like” structures (white arrows), predominantly in gp91phox KO lungs, morphologically resembling AFs. Higher magnification images are provided for improved visualization in the bottom of the figure.

**Figure 3.**
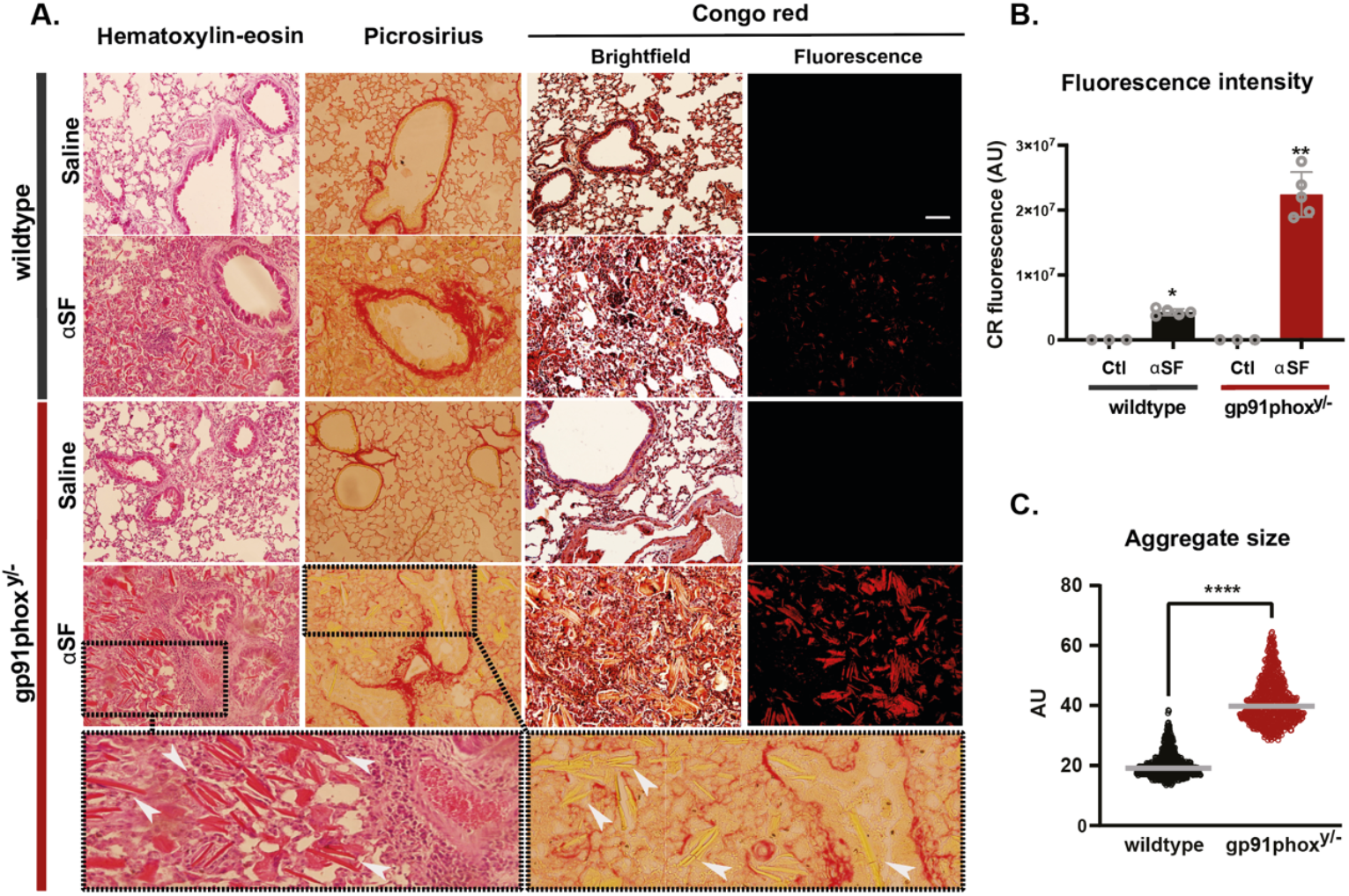
Characterization of lung injury induced by αSF instillation in WT and gp91phox KO mice. (**A**) Histological analysis of lung sections collected 4 h after αSF instillation, stained with hematoxylin and eosin (H&E) to evaluate tissue architecture, Picrosirius red for collagen deposition, and Congo red for amyloid detection. Lower panels show magnification of the areas highlighted in the rectangles, where it is possible to see (arrows) the presence of toothpick-like structure with fibrillar appearance in the lungs of gp91phox KO mice. (**B**) Quantification of Congo red fluorescence intensity, indicating the extent of amyloid deposition in lung tissue. (**C**) Measurement of aggregate size in lung sections from each experimental group. Data present in **panels B** and **C** are shown as mean ± SEM of 5 animals per condition. Statistical analysis was performed using one-way ANOVA in B: * = p < 0.05; ** = p < 0.01 (compared to the control group) and student’s t-test in C: **** = p < 0.01 compared to the control group. In these experiments, only male mice were used. Scale bar = 200 μm.

To confirm the amyloid nature of these structures, we performed Congo Red staining, which specifically labels β-sheet–rich aggregates. Quantification of Congo Red–positive structures showed a significantly greater number and larger average size of aggregates in gp91phox KO lungs compared to WT mice at 4 h post-instillation (**Figure 3B** and **C**). In WT lungs, only small aggregates were detected at this time point. Together, these findings suggest that αSFs are efficiently cleared from WT lungs within 4 h of instillation, whereas in gp91phox KO mice, fibrillar structures persist and accumulate, potentially exacerbating lung inflammation.

### αSF trigger NETs formation *in vivo* in a NOX-2-dependent manner

Having established that AFs induce NET formation in WT murine neutrophils *in vitro*, and that AF instilled into the lungs of WT and gp91phox KO mice recruit immune cells, including neutrophils, we next investigated whether AFs could trigger NET formation *in vivo*. For this, we stained αSF-instilled lungs with anti-H3Cit and anti-MPO antibodies, two NET associated proteins that, when colocalized, suggest NETosis. As shown in **Figure 4**, colocalization of H3Cit and MPO was observed exclusively in the lungs of WT mice instilled with αSF, confirming NET formation in response to αSF instillation. We did not notice the presence of NET markers in the lungs of αSF-instilled gp91phox KO mice, which is expected since these animals are ineffective in ROS-dependent NET production.

**Figure 4.**
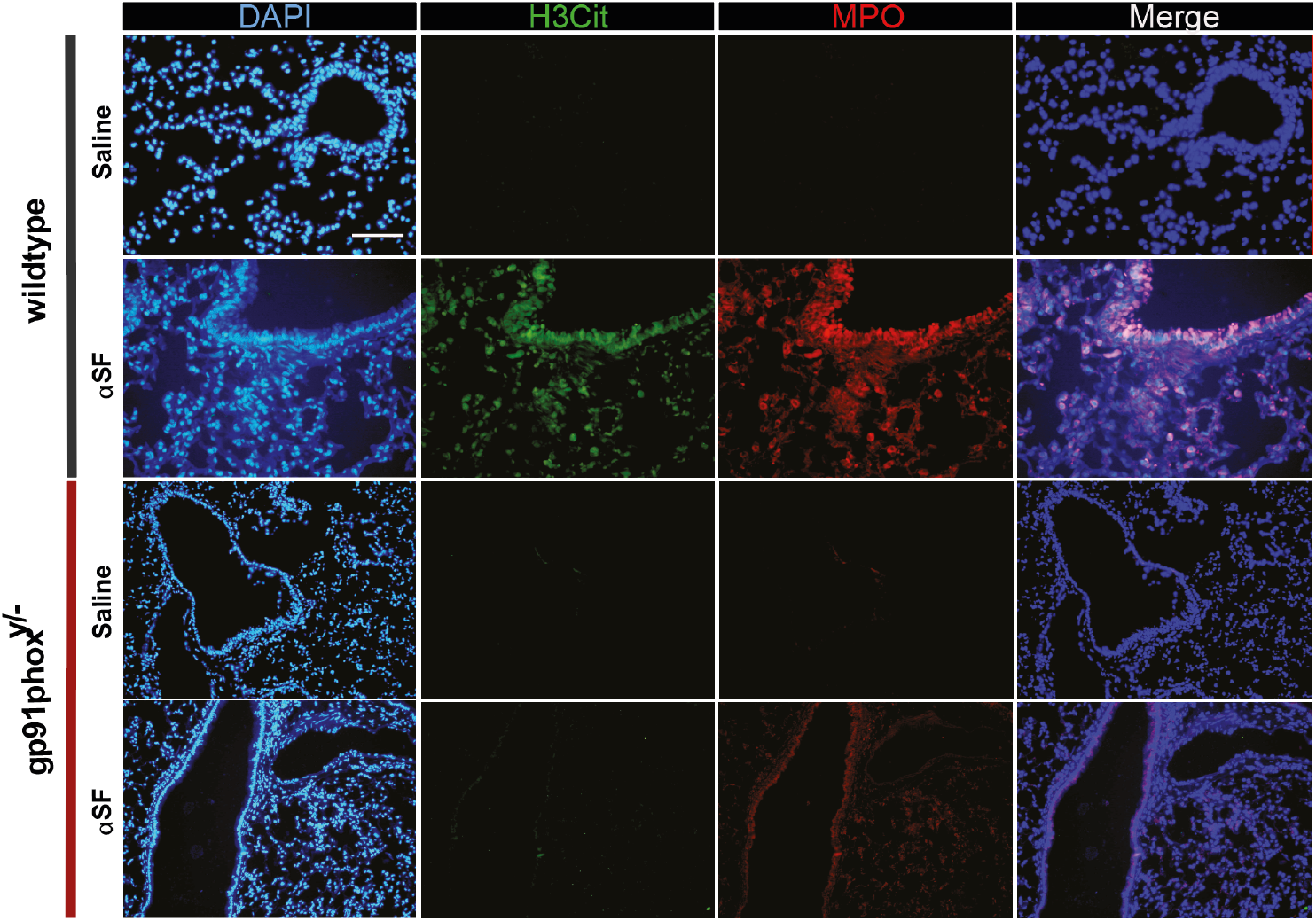
αSF induces NET formation *in vivo* in a NADPH oxidase 2-dependent manner. Immunofluorescence analysis of male lung sections of WT and gp91phox KO mice 4 h after intratracheal instillation with either saline or 50 μg of αSF. NETs were identified by co-localization of citrullinated histone H3 (green; anti-CitH3 antibody) and myeloperoxidase (red; anti-MPO antibody). Nuclei were counterstained with DAPI (blue). Scale bar = 200 μm.

### NETs mediate lung recovery after amyloid fibril instillation

To determine whether αSF deposition affects lung function, we evaluated lung pulmonary parameters both in male and female of WT and gp91phox KO mice 8 h after αSF instillation. For this, resistance and elastance were evaluated using invasive whole-body plethysmography, following exposure to increasing doses of the bronchoconstrictor agent methacholine. As shown in **Figure 5**, only the female and male of gp91phox KO mice instilled with αSF exhibited a significant increase in lung elastance, beginning at 9 mg/mL and 27 mg/mL of methacholine, respectively (red bars in **Figure 5A** and **C**). Notably, except for male gp91phox KO mice instilled with αSF and treated with 127 mg/mL of methacholine, no significant changes in lung resistance were observed in WT or KO animals, female or male, following αSF instillation, even when high doses of methacholine were used (**Figure 5B** and **D**). This selective increase in elastance - without a corresponding change in resistance - suggests that the damage is primarily localized to the lung parenchyma, without airway obstruction. In contrast, male and female WT mice 8 h after AF instillation did not display significant alterations in either lung elastance or resistance, even at higher methacholine concentrations (**Figure 5A-D**).

**Figure 5.**
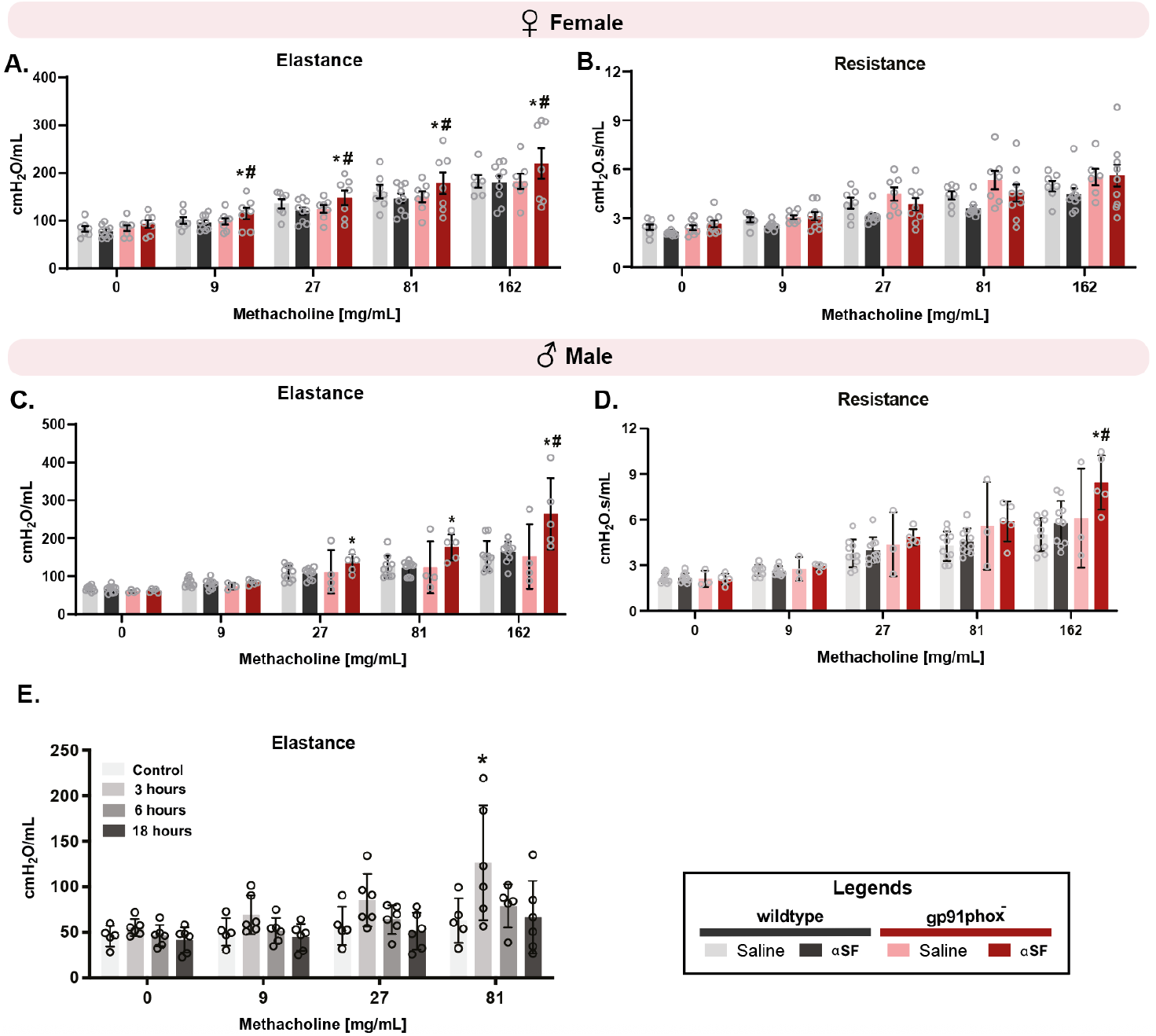
αSF triggers alterations in pulmonary function of mice. Lung injury was assessed by measuring elastance (**A** and **C**) and resistance (**B** and **D**) 8 h after instillation of 50 μg of AF, following increasing doses of the bronchoconstrictor agent methacholine. Results are presented separately by sex (**A** and **B**, female; **C** and **D**, male). (**E**) Elastance measurements with male WT mice instilled with AF as a function of time. Note that at 3h elastance if compromised in these lungs. Results expressed as mean ± SEM. * p < 0.05 compared to the control group instilled with saline (Ctl); # p < 0.05 compared to the WT group instilled with αSF; Female: n = 8 for Ctl and 9 for instilled with αSF in both groups. Male: n = 10 for WT in both groups, 3 for gp91phox^y/-^ Ctl and 6 for gp91phox^y/-^ αSF. Kinetic data with males: n = 5 Control.

Since αSFs were no longer detectable in the lungs of WT mice 8 h after instillation - the time point at which impaired lung function was observed in KO animals - we evaluated lung elastance in male WT mice at earlier time points (**Figure 5E**). Remarkably, a significant impairment in lung function was already evident 3 h after αSF instillation in WT mice, with progressive recovery over time. This finding suggests that αSFs cause a transient disruption of lung mechanics in WT animals, which is resolved as the fibrils are effectively cleared.

### Are NET’s components able to digest AFs into small species?

Since NETs are composed of various proteases, including elastase, next we investigated whether the proteases associated with NETs could degrade αSF into smaller species *in vitro*. To this end, αSF was incubated with SPAN (Supernatant of PMA-Activated human Neutrophils), a suspension enriched in NETs or with purified elastase (**Figure 6**).

**Figure 6:**
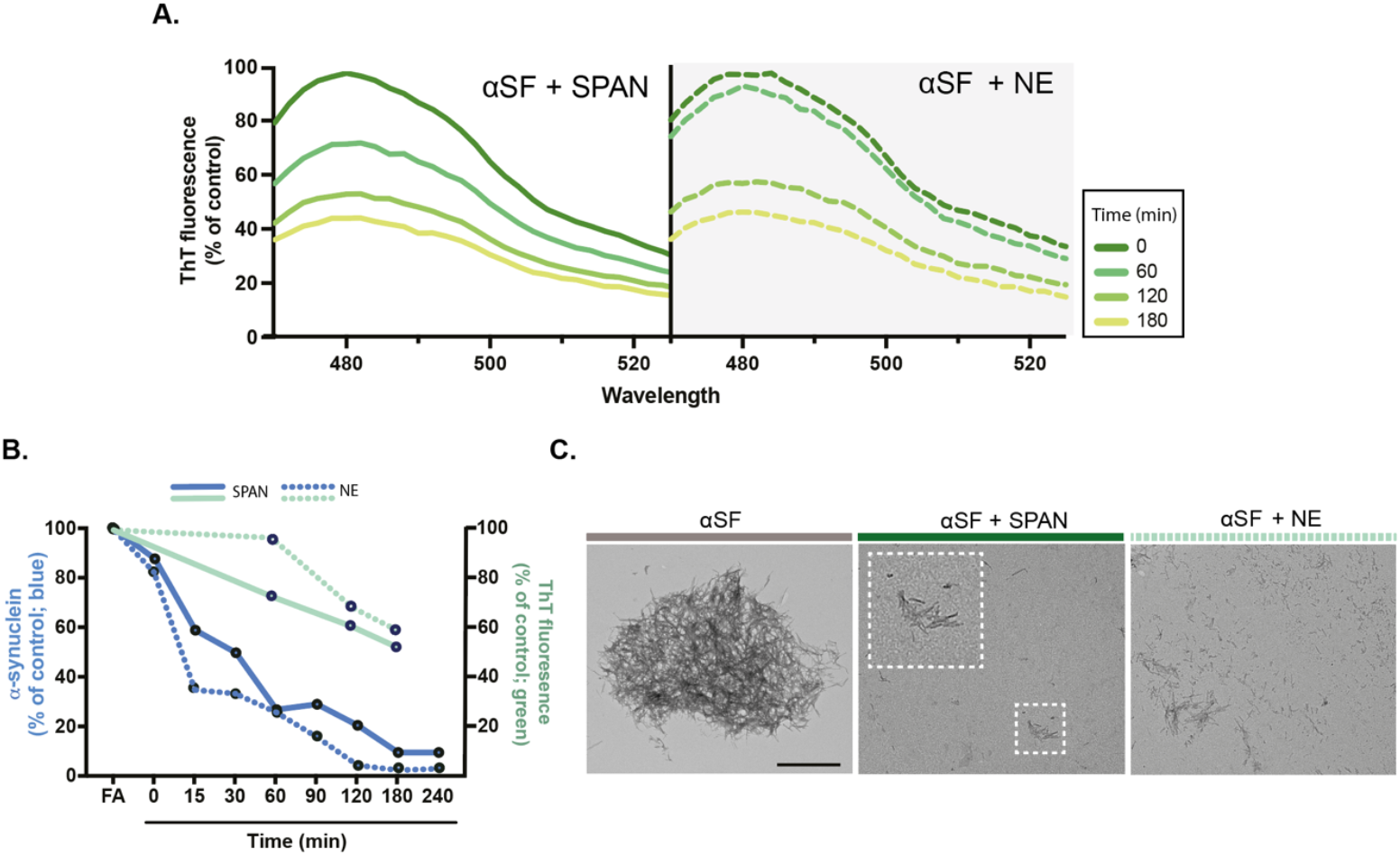
SPAN and Elastase digest αSF into small fragments. (**A**) Thioflavin T (ThT) fluorescence emission spectra of aSF at different times in the absence (**left panel, control**) and in the presence of SPAN (**middle**) or Elastase (**right**). (**B**) Quantification of the ThT emission spectra extracted from **panel A** (green): SPAN (**continuous line**) and Elastase (**dashed line**). The intensities of αSyn monomers bands, quantified from the western blots presented in **Supp Figure 2**, are displayed in blue (SPAN, continuous line and Elastase, dashed line). (**C**) TEM images for the mature aSF (**left panel**), fragments generated after 2h in the presence of SPAN (**middle panel**) and Elastase (**right panel**); Scale bar = 500 nm. ThT emission - Ex = 450 nm, Em = 482. [SPAN] = 1 μg/μL; [Elastase] = 10 μg/μL; [aSF] = 5 μM.

As shown in **Figures 6A**, Thioflavin T (ThT) fluorescence - an indicator of amyloid fibrils - decreased in a time-dependent manner upon incubation of αSF with SPAN (middle panel) or elastase (right panel), indicating progressive fibril degradation while untreated αSF maintained stable ThT fluorescence over time (not shown). Transmission electron microscopy (TEM) images (**Figure 6C**) further confirmed the structural disruption of αSF after 2 h of incubation with SPAN or elastase (middle and right panels, respectively), revealing the transformation from intact fibrils to fragmented structures. Western blot analysis using an anti-α-synuclein antibody (Supplementary Figure 2) showed a reduction in the monomeric α-synuclein band, consistent with proteolytic processing by SPAN and elastase, and their quantification together with ThT changes are displayed in **Figure 6B**.

## Discussion

Since the seminal description of NETs by Brinkmann and coauthors (2004), these structures have been implicated in a wide range of pathological conditions^20^. Our group was among the first to propose a direct link between NET formation and neurodegenerative diseases caused by amyloid deposition^27^. Specifically, we demonstrated that amyloid fibrils of various compositions, including α-synuclein, can induce NET release from human neutrophils *in vitro*. Not only were AFs able to trigger NETosis, but they were also digested by proteases, mainly elastase, embedded within these traps into toxic oligomeric species. We reported the simultaneous presence of AFs and NET markers in human tissues, specifically in lung and skin biopsies from patients who died from primary systemic amyloidosis^27^.

However, despite this striking observation, it remained unclear whether amyloid fibrils could actively recruit neutrophils or directly induce NET release *in vivo* in an animal model. To address this, we employed a mouse model deficient in NOX-2 activity, a key enzyme required for the generation of ROS in phagocytes, mainly neutrophils. These mice, originally developed as a model for chronic granulomatous disease^29^, are unable to produce ROS and, consequently, fail to generate ROS-dependent NETs, as demonstrated in this study (**Figure 1**). This makes them a valuable tool to investigate the contribution of NETs in pathologies where ROS-mediated mechanisms drive NET formation.

Consistent with our *in vitro* observations in human neutrophils, we found that α-synuclein fibrils (αSF) stimulated NET formation in murine neutrophils in a ROS-dependent manner (**Figure 1A**). This response was evident in WT mice but absent in gp91phox-deficient animals, as confirmed by NET formation kinetics measured through Sytox Green fluorescence and supporting fluorescence images (**Figure 1B**). Supplementing gp91phox-deficient neutrophils with low concentrations of hydrogen peroxide (1 μM H_2_O_2_) restored their ability to release NETs upon αSF stimulation, reaching levels comparable to WT controls (**Figure 1C** and **D**). Previous studies have shown that modulation of the neutrophil microenvironment, such as extracellular pH, can spontaneously induce NET release^33^. In our experiments, the low dose of H_2_O_2_ used did not induce NETs by itself, in contrast to higher concentrations previously reported to do so^33^. Together, these findings strongly suggest that the impaired NET formation in gp91phox KO neutrophils results from ROS deficiency rather than from intrinsic defects in the molecular machinery regulating NETosis.

Once we confirmed that αSF–induced NETosis occurs *in vitro* in murine neutrophils, we next investigated whether αSF could recruit neutrophils *in vivo*. To this end, 50 μg of αSF were instilled intratracheally into the lungs of both WT and gp91phox-KO mice of both sexes. Eight hours later, leukocyte recruitment was observed in both sexes (**Figure 2A** and **G**), with neutrophils representing a major proportion of the infiltrating cells (**Figure 2 C** and **I**). Notably, gp91phox-KO mice of both sexes recruited significantly more leukocytes than WT controls, likely as a compensatory response to their ROS deficiency (**Figure 2A, C, G**, and **I**). This enhanced neutrophil influx was accompanied by increased MPO activity (**Figure 2D** and **J**). Elevated blood levels of MPO and other neutrophil-derived enzymes have previously been linked to cognitive decline in AD^34^, underscoring a possible pathological relevance of dysregulated neutrophil activity in neurodegeneration.

Sex-specific differences were also observed. Male WT mice recruited mononuclear cells at levels comparable to KO animals, whereas female WT mice displayed no detectable mononuclear cell recruitment following αSF instillation. Although the basis of this difference remains unclear, similar sex-dependent immune responses have been reported^35,36^. Furthermore, male KO mice recruited more immune cells than their female counterparts, possibly reflecting their hemizygous status for the X-linked gp91phox gene. A comparable phenomenon was described by Pollock et al. (1995)^29^, who first generated these mice and reported enhanced immune cell recruitment - including neutrophils - into the peritoneal cavity after thioglycolate challenge compared to WT controls.

Lung sections from both WT and gp91phox KO mice, examined 4 h after αSF instillation, showed intense cellular infiltration (H&E) and alveolar obstruction by exogenous AFs (confirmed by CR staining) (**Figure 3B** and **C**). These alterations were more pronounced in gp91phox KO animals than in WT (**Figure 3A**). In addition to cell infiltration, both genotypes displayed increased collagen deposition within the parenchyma, a process known to represent an early hallmark of acute pulmonary inflammation triggered by antigenic stimulation or injury^37^.

The colocalization of H3Cit-a hallmark post-translational modification of NETs^38^-with MPO confirmed NET formation *in vivo* in WT mice. In contrast, H3Cit was undetectable in gp91phox-KO mice, either alone or in colocalization with MPO, indicating that ROS generation is an upstream requirement for amyloid fibril–induced NET release (**Figure 4**), consistent with findings for other NET stimuli^18,24^. The presence of MPO staining in gp91phox-KO mice without detectable H3Cit further supports the conclusion that αSF promote neutrophil recruitment but cannot trigger NET formation in the absence of functional NOX-2 (**Figure 1**). Collectively, these findings demonstrate that αSF recruit neutrophils and induce NETosis *in vivo* in a NOX-2-dependent manner.

As previously demonstrated in our *in vitro* studies with transthyretin fibrils^27^, NET-associated proteases - particularly neutrophil elastase - can degrade amyloid fibrils into smaller, toxic species. In line with this, 4 h after αSF instillation, large toothpick-like structures were still evident in the lungs of gp91phox-deficient mice (**Figure 3A**), whereas WT animals displayed fewer and smaller Congo red–positive amyloid-like deposits, suggesting a possible role of NETs in amyloid clearance.

Given that our earlier work had focused on TTR fibrils, we next examined whether NET-associated proteases could similarly digest αSF. For this, αSF was incubated *in vitro* with SPAN or neutrophil elastase for 2 h, and fibril degradation was monitored by western blotting (**Supp. Fig. 2)** and Thioflavin-T (ThT) fluorescence (**Figure 6**). The progressive disappearance of the αSF-specific band, together with a time-dependent decrease in ThT fluorescence, indicated amyloidolysis and loss of amyloid structure. Transmission electron microscopy further confirmed this breakdown, revealing that SPAN- and neutrophilic elastase-mediated digestion produced shorter and less clustered fibrils. Taken together, these results strongly suggest that, like TTR, αSF can be digested by NET-associated proteases into smaller, potentially toxic species.

To evaluate the impact of aggregates and the NETs induced by them in the lung, we performed functional assays using plethysmography to measure elastance and resistance. Our analysis revealed no evidence of pulmonary dysfunction 8 h after αSF instillation in the WT mice (**Figure 5A-D**). However, a transient reduction in lung elastance was detected at earlier time points (**Figure 5E**), reflecting inflammation at the level of the lung parenchyma. Regarding the gp91phox-KO mice, even 8 h after αSF instillation, lung elastance was still altered. These results indicate that the presence of αSF causes acute lung injury and that their efficient elimination in WT mice protects the lungs from sustained damage, the persistence of αSF in the KO mice, which results in lung injury. Whether these smaller species can disseminate to other organs and tissues and seed pathology remains to be determined.

The involvement of neutrophils in the prognosis of amyloidosis has been previously reported. For instance, Zenaro and colleagues (2015)^39^ identified neutrophil infiltration in the brain parenchyma of two transgenic mouse models of Alzheimer’s disease (5xFAD and 3xTg-AD). In addition to the presence of neutrophils, the authors detected NET-associated markers within the brain tissue. Depletion of neutrophils in these models significantly restored cognitive function, and ameliorated disease-related symptoms. In another study using the APP/PS1 mouse model of AD^40^, researchers reported elevated levels of neutrophils in the brain compared to wild-type controls. Remarkably, inhibiting neutrophil migration from the bloodstream into the brain parenchyma rescued cognitive deficits, further implicating neutrophils in the pathogenesis of neurodegenerative diseases^39,40^. However, it remains unclear whether the observed improvement was due to the absence of neutrophils themselves or the lack of NET production.

More recently, Hancock and coworkers (2024)^41^ analyzed how neutrophils may contribute to the clearance of human AL amyloid extracts, using murine amyloidoma models composed of ALλ, which is refractory to clearance, and ALκ, which is readily cleared in mice. Their data showed that more neutrophils were recruited to ALκ amyloid masses as compared to the ALλ material, which was devoid of neutrophils. Interestingly their data showed by *ex vivo* analyses that neutrophils do not efficiently phagocytose amyloids directly but, instead, in the ALκ amyloidoma there was an abundant presence of NETs, which were absent in the ALλ amyloidomas. These findings are in accordance with ours, which suggest the participation of NETs in amyloid clearance.

As we recently, showed, NET is a double-edge sword^42^. In one hand, it can, as shown here, degrade AF sparing the lungs or any other tissue from the detrimental presence of amyloid fibrils. On the other, NET presence can be detrimental to the surrounding tissue, where it is formed. Regarding the peripheral nervous system, our group showed that MPO and histones present in NET structures were very toxic to neurons and Schwann cells^42^.

Beyond the data presented here supporting a role for neutrophils and their extracellular traps in amyloidosis, it is important to acknowledge certain limitations of the model employed. To the best of our knowledge, α-synuclein aggregates have not been detected in the lungs of patients with Parkinson’s disease. Our choice of the lung as a model for amyloid clearance was guided by several considerations: (i) pulmonary involvement is relatively common in systemic amyloidosis, although it rarely manifests with overt clinical symptoms^43^; (ii) the lung is a well-established model for investigating acute inflammation^44^; (iii) its compartment provides direct access for immune recruitment and interactions; and (iv) the lung–brain axis has recently emerged as an area of growing interest, with increasing evidence that these organs may communicate through neuroanatomical, endocrine, and immunological pathways^45–49^.

In summary (**Figure 7**), this study demonstrates that the instillation of amyloid fibrils in the lungs triggers neutrophil recruitment and the formation of neutrophil extracellular traps *in vivo* through a reactive oxygen species–dependent pathway. These traps contribute to the clearance of amyloid fibrils and promote the recovery of pulmonary function by degrading the amyloid fibrils. Taken together, our findings highlight neutrophils as active participants in amyloid pathology and point to their potential role in shaping new therapeutic strategies across amyloid-related diseases.

**Figure 7.**
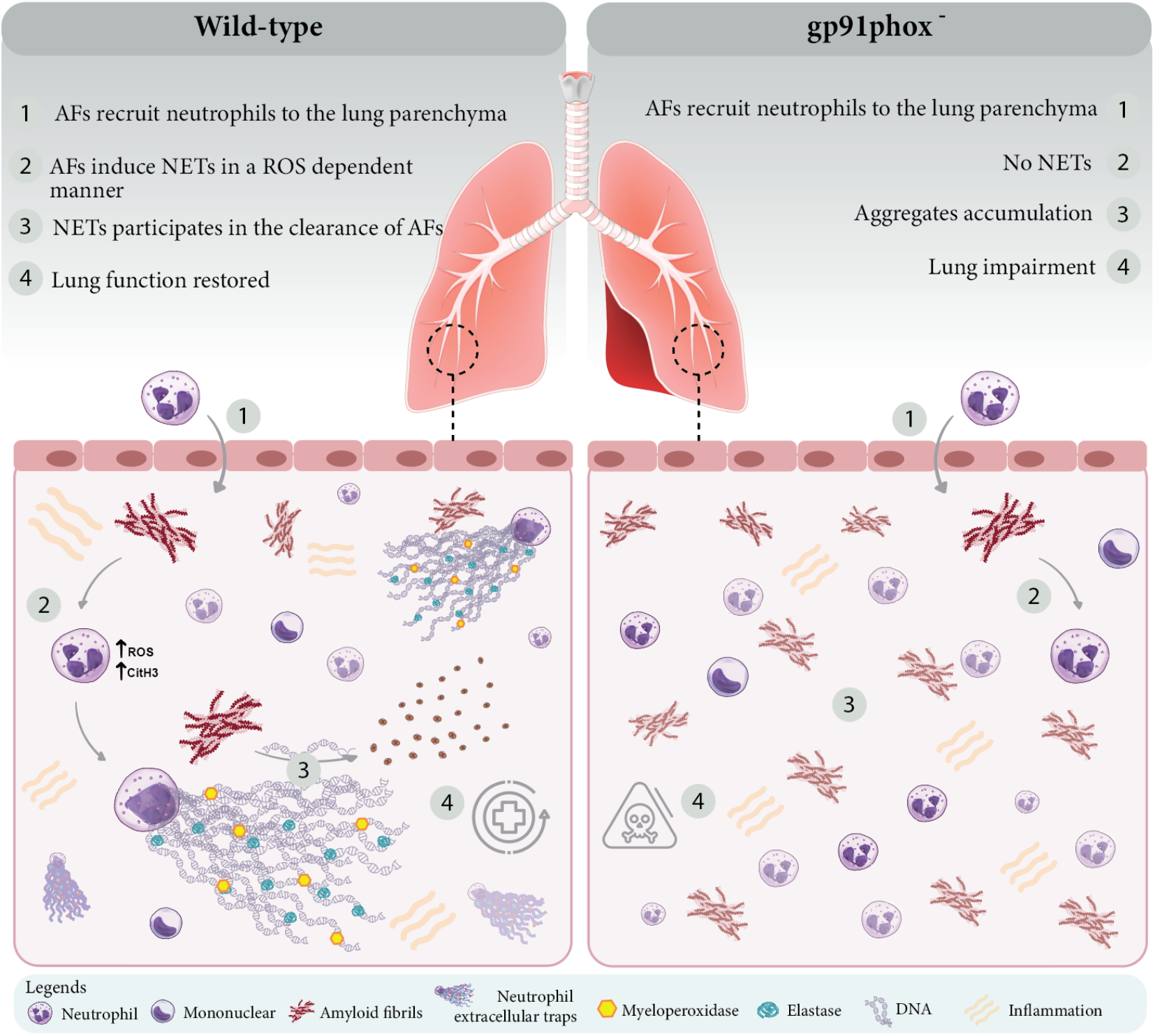
Summary of the main findings. Amyloid fibrils are deposit in the lungs of WT and KO animals causing mal functioning. (**1**) Neutrophils, as a first line of defense, are recruited to the deposition site in WT or KO animals causing acute inflammation. (**2**) Only in WT animals, AF induce NET formation in a ROS-dependent manner. (**3**) The NET-components, mainly elastase, degrade AF into small fragments, what ameliorate lung functioning (**4**), what does not take place in KO mice, where AF persist.

## Notes

### Competing Interest Statement

The authors have declared no competing interest.

## References

1. Tyedmers, J., Mogk, A. & Bukau, B. Cellular strategies for controlling protein aggregation. Nat. Rev. Mol. Cell Biol. 11, 777–788 (2010).

2. Bertram, L. & Tanzi, R. E. The genetic epidemiology of neurodegenerative disease. J. Clin. Invest. 115, 1449–1457 (2005).

3. Fernandes, L., Cardim-Pires, T. R., Foguel, D. & Palhano, F. L. Green Tea Polyphenol Epigallocatechin-Gallate in Amyloid Aggregation and Neurodegenerative Diseases. vol. 15 (2021).

4. Louros, N., Schymkowitz, J. & Rousseau, F. Mechanisms and pathology of protein misfolding and aggregation. Nat. Rev. Mol. Cell Biol. (2023) doi:10.1038/s41580-023-00647-2.

5. Wilson, M. R., Satapathy, S. & Vendruscolo, M. Extracellular protein homeostasis in neurodegenerative diseases. Nature Reviews Neurology (2023) doi:10.1038/s41582-023-00786-2.

6. Rinauro, D. J., Chiti, F., Vendruscolo, M. & Limbocker, R. Misfolded protein oligomers: mechanisms of formation, cytotoxic effects, and pharmacological approaches against protein misfolding diseases. Molecular Neurodegeneration vol. 19 Preprint at 10.1186/s13024-023-00651-2 (2024).

7. Oliveira, J. T. et al. Oligomeric α-Synuclein induces skin degeneration in reconstructed human epidermis. Neurobiol. Aging 113, 108–117 (2022).

8. Breydo, L. & Uversky, V. N. Structural, morphological, and functional diversity of amyloid oligomers. FEBS Lett. 589, 2640–2648 (2015).

9. Blum-Degena, D. et al. Interleukin-1β and interleukin-6 are elevated in the cerebrospinal fluid of Alzheimer’s and de novo Parkinson’s disease patients. Neurosci. Lett. 202, 17–20 (1995).

10. Paganelli, R. et al. Proinflammatory cytokines in sera of elderly patients with dementia: levels in vascular injury are higher than those of mild-moderate Alzheimer’s disease patients. Exp. Gerontol. 37, 257–263 (2002).

11. Balkhi, S., Di Spirito, A., Poggi, A. & Mortara, L. Immune modulation in Alzheimer’s disease: From pathogenesis to immunotherapy. Cells 14, (2025).

12. Williams, G. P. et al. CD4 T cells mediate brain inflammation and neurodegeneration in a mouse model of Parkinson’s disease. Brain 144, 2047–2059 (2021).

13. Lawrence, S. M., Corriden, R. & Nizet, V. How neutrophils meet their end. Trends Immunol. 41, 531–544 (2020).

14. Zhang, F. et al. Neutrophil diversity and function in health and disease. Signal Transduct. Target. Ther. 9, 343 (2024).

15. Amulic, B., Cazalet, C., Hayes, G. L., Metzler, K. D. & Zychlinsky, A. Neutrophil function: from mechanisms to disease. Annu. Rev. Immunol. 30, 459–489 (2012).

16. Brinkmann, V. et al. Neutrophil Extracellular Traps Kill Bacteria. Science 303, 1532–1535 (2004).

17. Kenny, E. F. et al. Diverse stimuli engage different neutrophil extracellular trap pathways. Elife 6, 1–21 (2017).

18. Petretto, A. et al. Neutrophil extracellular traps (NET) induced by different stimuli: A comparative proteomic analysis. PLoS One 14, e0218946 (2019).

19. Neumann, A., Brogden, G. & von Köckritz-Blickwede, M. Extracellular traps: An ancient weapon of multiple kingdoms. Biology (Basel) 9, 34 (2020).

20. Papayannopoulos, V. Neutrophil extracellular traps in immunity and disease. Nat. Rev. Immunol. 18, 134–147 (2018).

21. Zhang, S., Guo, M., Liu, Q., Liu, J. & Cui, Y. Neutrophil extracellular traps induce a hypercoagulable state in glioma. Immun. Inflamm. Dis. 9, 1383–1393 (2021).

22. Zhang, S., Guo, M., Liu, Q., Liu, J. & Cui, Y. Neutrophil extracellular traps induce thrombogenicity in severe carotid stenosis. Immun. Inflamm. Dis. 9, 1025–1036 (2021).

23. Stoiber, W., Obermayer, A., Steinbacher, P. & Krautgartner, W.-D. The role of reactive oxygen species (ROS) in the formation of extracellular traps (ETs) in humans. Biomolecules 5, 702–723 (2015).

24. Parker, H., Dragunow, M., Hampton, M. B., Kettle, A. J. & Winterbourn, C. C. Requirements for NADPH oxidase and myeloperoxidase in neutrophil extracellular trap formation differ depending on the stimulus. Journal of Leukocyte Biology 92, 841–849 (2012).

25. Neubert, E., Meyer, D., Kruss, S. & Erpenbeck, L. The power from within - understanding the driving forces of neutrophil extracellular trap formation. J. Cell Sci. 133, jcs241075 (2020).

26. Pieterse, E., Rother, N., Yanginlar, C., Hilbrands, L. B. & van der Vlag, J. Neutrophils discriminate between lipopolysaccharides of different bacterial sources and selectively release neutrophil extracellular traps. Front. Immunol. 7, 1–13 (2016).

27. Azevedo, E. P. C. et al. Amyloid fibrils trigger the release of neutrophil extracellular traps (NETs), causing fibril fragmentation by NET-associated elastase. J. Biol. Chem. 287, 37206–37218 (2012).

28. Azevedo, E. P. et al. A metabolic shift toward pentose phosphate pathway is necessary for amyloid fibril- and phorbol 12-myristate 13-acetate-induced neutrophil extracellular trap (NET) formation. J. Biol. Chem. 290, 22174–22183 (2015).

29. Pollock, J. D. et al. Mouse model of X-linked chronic granulomatous disease, an inherited defect in phagocyte superoxide production. Nat. Genet. 9, 202–209 (1995).

30. Ermert, D. et al. Mouse neutrophil extracellular traps in microbial infections. J. Innate Immun. 1, 181–193 (2009).

31. Coelho-Cerqueira, E., Carmo-Gonçalves, P., Pinheiro, A. S., Cortines, J. & Follmer, C. α-Synuclein as an intrinsically disordered monomer--fact or artefact? FEBS J. 280, 4915–4927 (2013).

32. Ferreira, T. P. T. et al. Intranasal flunisolide suppresses pathological alterations caused by silica particles in the lungs of mice. Front. Endocrinol. (Lausanne) 11, 388 (2020).

33. Behnen, M., Möller, S., Brozek, A., Klinger, M. & Laskay, T. Extracellular acidification inhibits the ROS-dependent formation of neutrophil extracellular traps. Front. Immunol. 8, 184 (2017).

34. Wu, C.-Y. et al. Neutrophil activation in Alzheimer’s disease and mild cognitive impairment: A systematic review and meta-analysis of protein markers in blood and cerebrospinal fluid. Ageing Res. Rev. 62, 101130 (2020).

35. Klein, S. L. & Flanagan, K. L. Sex differences in immune responses. Nat. Rev. Immunol. 16, 626–638 (2016).

36. Lingappan, K. et al. Sex-specific differences in hyperoxic lung injury in mice: implications for acute and chronic lung disease in humans. Toxicol. Appl. Pharmacol. 272, 281–290 (2013).

37. Sousse, L. E. et al. Long-term pulmonary dysfunction and collagen deposition after burn and smoke inhalation is mediated by reactive oxygen species, asymmetric dimethylarginine, and arginase. in D35. OXIDANTS AND ANTIOXIDANTS (American Thoracic Society, 2011). doi:10.1164/ajrccm-conference.2011.183.1_meetingabstracts.a5958.

38. Hamam, H. J. & Palaniyar, N. Post-translational modifications in NETosis and NETs-mediated diseases. Biomolecules 9, 369 (2019).

39. Zenaro, E. et al. Neutrophils promote Alzheimer’s disease-like pathology and cognitive decline via LFA-1 integrin. Nat. Med. 21, 880–886 (2015).

40. Qi, F. et al. VEGF-A in serum protects against memory impairment in APP/PS1 transgenic mice by blocking neutrophil infiltration. Mol. Psychiatry 28, 4374–4389 (2023).

41. Hancock, T. J. et al. Neutrophils enhance the clearance of systemic amyloid deposits in a murine amyloidoma model. Front. Immunol. 15, 1487250 (2024).

42. de Siqueira Santos, R. et al. Peripheral nervous system is injured by neutrophil extracellular traps (NETs) elicited by nonstructural (NS) protein-1 from Zika virus. FASEB J. 37, e23126 (2023).

43. Milani, P. et al. The lung in amyloidosis. Eur. Respir. Rev. 26, 170046 (2017).

44. Potey, P. M., Rossi, A. G., Lucas, C. D. & Dorward, D. A. Neutrophils in the initiation and resolution of acute pulmonary inflammation: understanding biological function and therapeutic potential. J. Pathol. 247, 672–685 (2019).

45. Hosang, L. et al. The lung microbiome regulates brain autoimmunity. Nature 603, 138–144 (2022).

46. Ziaka, M. & Exadaktylos, A. Gut-derived immune cells and the gut-lung axis in ARDS. Crit. Care 28, 220 (2024).

47. Bajinka, O., Simbilyabo, L., Tan, Y., Jabang, J. & Saleem, S. A. Lung-brain axis. Crit. Rev. Microbiol. 48, 257–269 (2022).

48. Li, Y. et al. Revealing a causal relationship between gut microbiota and lung cancer: a Mendelian randomization study. Front. Cell. Infect. Microbiol. 13, 1200299 (2023).

49. Wang, L. et al. The bidirectional gut-lung axis in chronic obstructive pulmonary disease. Am. J. Respir. Crit. Care Med. 207, 1145–1160 (2023).

