## Supplemental figures for "Neutrophil Extracellular Traps Mediate *In Vitro* and *In Vivo* Degradation of α-Synuclein Amyloid Fibrils"

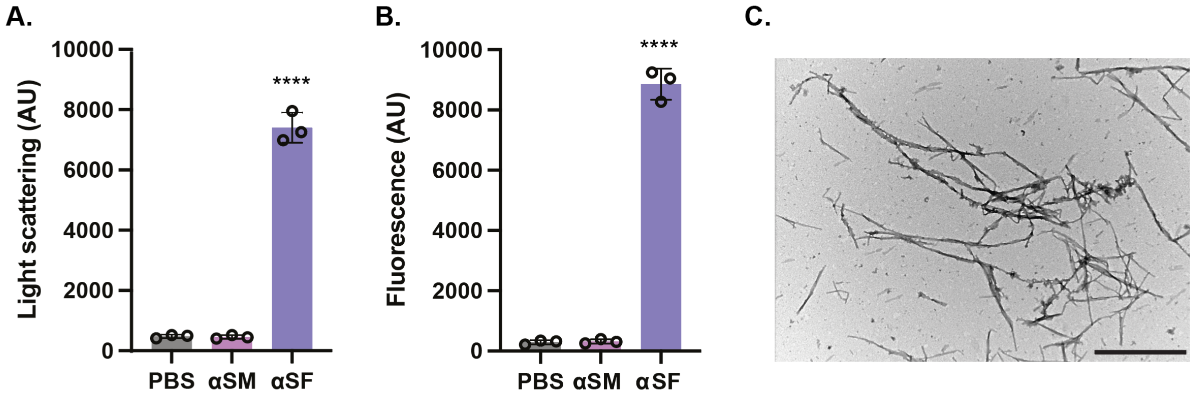


**Supplementary Figure 1 – Characterization of the α-synuclein amyloid fibrils used in the study.** Following purification and aggregation, α-synuclein amyloid fibrils were characterized by light scattering at 330 nm (**A**) and thioflavin T binding assay at 482 nm (**B**). Fibril morphology was visualized by transmission electron microscopy (TEM) (**C**). αSM = α-synuclein monomers; αSF = α-synuclein fibrils. **** $P < 0.0001$. Scale bars: 1 μm.

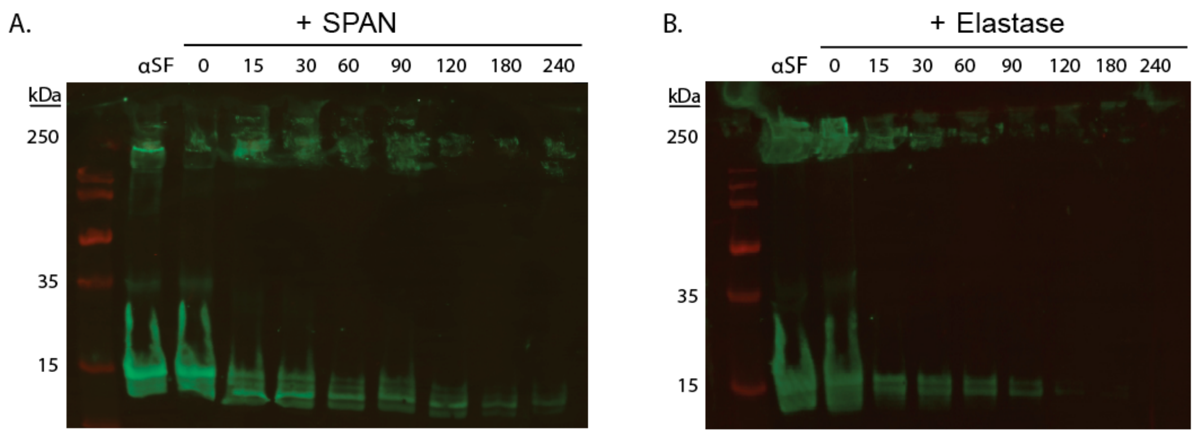


**Supplementary Figure 2 - Amyloid fibrils are digested into smaller species when incubated with SPAN or Elastase.** (**A**) Representative kinetics of αSF (5 µM) digestion by SPAN (1 µg/mL) or (**B**) Elastase (10 µg/mL) are shown in a Western Blotting using anti-αSF antibody at 0, 15, 30, 60, 120, 180, and 240 min. Note that over time, antibody labeling decreases, indicating digestion of these fibrils.
